# Local Bilayer Hydrophobicity Modulates Membrane Protein Stability

**DOI:** 10.1101/2020.09.01.277897

**Authors:** Dagan C. Marx, Karen G. Fleming

## Abstract

Through the insertion of nonpolar side chains into the bilayer, the hydrophobic effect has long been accepted as a driving force for membrane protein folding. However, how the changing chemical composition of the bilayer affects the magnitude side chain transfer free energies 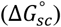 has historically not been well understood. A particularly challenging region for experimental interrogation is the bilayer interfacial region that is characterized by a steep polarity gradient. In this study we have determined the 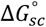 for nonpolar side chains as a function of bilayer position using a combination of experiment and simulation. We discovered an empirical correlation between the surface area of nonpolar side chain, the transfer free energies, and the local water concentration in the membrane that allows for 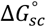 to be accurately estimated at any location in the bilayer. Using these water-to-bilayer 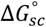 values, we calculated the interface-to-bilayer transfer free energy 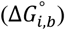. We find that the 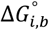 are similar to the “biological”, translocon-based transfer free energies, indicating that the translocon energetically mimics the bilayer interface. Together these findings can be applied to increase the accuracy of computational workflows used to identify and design membrane proteins, as well as bring greater insight into our understanding of how disease-causing mutations affect membrane protein folding and function.

## Introduction

The cellular functions of a protein are regulated by the intrinsic thermodynamic stability of the three-dimensional structure adopted by its polypeptide chain.^1–4^ The stabilities of many water-soluble proteins have been experimentally measured *in vitro*, resulting in an understanding of the forces that regulate structure and function of this class of proteins.^5^ For membrane proteins, which make up approximately one third of the eukaryotic proteome,^6^ clarity on the balance of forces that determine their thermodynamic stabilities is much less mature.^7^ This lag in knowledge derives from the fact that membrane proteins are aggregation prone in aqueous solutions and must embed themselves into a hydrophobic environment such as detergent micelles or phospholipid bilayers to fold and function.^8–10^ The aggregation propensity renders traditional experimental approaches for measuring protein stabilities untenable for most membrane proteins.^7^

The stabilities of membrane proteins are directly linked to the energetics of burying amino acid side chains into the phospholipid bilayer, also referred to as the side chain transfer free energy change 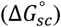.^7,10–12^ Both the magnitude and sign of the water-to-bilayer 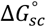 for a side chain in a transmembrane domain (TMD) depend on the exact chemical compositions of the two end-point solvents (water and bilayer).^13–22^ There is the additional complexity in that a bilayer is not a simple endpoint solvent. Biological membranes contain two chemically distinct regions: the largely desolvated hydrocarbon core and the chemically-heterogeneous interface.^23,24^ Along the *z*-coordinate of the bilayer normal, each interface is an approximately 15 Å wide region separating bulk water from the dehydrated center, and the transition between these two endpoints is characterized by a continuously changing water concentration.^15,24^ This water gradient should result in a position-dependence on the free energies of transfer for side chains along the interface. Arguably, these membrane interfaces represent half of the bilayer volume, and a full understanding of membrane protein folding forces depends on the energetics imposed by this steeply changing polarity of this region.

The hydrophobic effect has empirically been related to the (water) solvent accessible surface area (ASA) of the side chain through an energy termed the nonpolar solvation parameter, *σ_NP_*.^16,18,19,25–28^ This relationship quantifies the energy (cal mol^−1^) gained per Å^2^ of nonpolar surface area removed from water and buried in a nonpolar solvent. Across a wide range of model-protein systems and membrane mimics, the *σ_NP_* value shows a remarkably precise value, ranging from −23 to −25 cal mol^−1^ Å^−2^.^16,18,19^ In the vast majority of studies, this singular *σ_NP_* has traditionally been the basis for estimating side chain-bilayer partitioning as well as for identifying polypeptide sequence stretches that are likely to adopt transmembrane location. An exception to this is a single water-to-bilayer-interface study that found a *σ_NP_* value for that reaction equal to −12 cal mol^−1^ Å^−2^.

One challenge with all of these studies is a precise mapping of the *z*-position in the membrane to which each *σ_NP_* corresponds.^17^ Moreover, how to relate partitioning energies to a bilayer location is arguably impossible to know for those studies employing organic solvents as the bilayer mimic. Even with these drawbacks, usage of the nonpolar solvation parameter and accompanying transfer free energy changes for nonpolar side chains have proven to be very powerful for identification of transmembrane segments from sequence.

Nevertheless, these energy values are still a somewhat crude tool for estimating the stabilities of these polypeptide regions at different locations in the chemically-heterogeneous bilayer. Here we take advantage of the naturally-occurring water gradient in the bilayer to ask how this changing water concentration affects the energetics of nonpolar side chains and accordingly, the nonpolar solvation parameter. We accomplish these measurements using a natively-folded protein that possesses a very favorable folding free energy and that is firmly anchored in its position within the experimental bilayer.^29^ Using this setup, we have resolved the bilayer *z*-position dependence of *σ_NP_* using a combination of protein folding titrations, thermodynamic cycles and molecular dynamics simulations.

We discovered a linear correlation between *σ_NP_* and the local water concentration across the bilayer normal. This relationship essentially quantifies the hydrophobic effect across the bilayer interface and enables prediction of the functional form for a depth-dependent solvation parameter, *σ_NP_* (*z*), and more generally, functions describing side-chain transfer free energies, 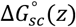, across the bilayer. These energy values should improve the accuracy of membrane protein structure prediction and design algorithms. Additionally, we use our findings to better understand the thermodynamic basis for how the Sec translocon aids α-helical membrane protein folding *in vivo*.

## Materials and Methods

### OmpLA Variant Expression, Purification, Activity Assay and Folding Titrations

The OmpLA variants were engineered, expressed, and purified from inclusion bodies as previously described.^15,16^ Each variant was verified to have enzymatic activity while folded into DLPC large unilamellar vesicles (LUV) using a previously developed colorimetric assay.^15,16^ Folding titrations were also set up and performed as previously described in detail.^15,16,18,30,31^ Briefly, unfolded OmpLA variants added to DLPC LUVs (diameter = 100nm) in 5 M guanidine hydrochloride (GdnHCl). After overnight incubation at 37 °C, samples were diluted to varying GdnHCl concentrations ranging from 1-5 M at 0.08 M increments (51 total samples). Following at least a 40-hour incubation in a rotating incubator at 37 °C, intrinsic tryptophan fluorescence was measured at 330 nm on a PC1 Fluorometer with cross-polarization (ISS, Champaign, IL, USA).

### Fitting OmpLA variant 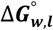 and σ_NP_

The resulting titration data were fit to a three-state linear extrapolation model to extract the stability 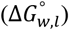 of each variant using previously determined *m*-values using the Igor Pro 8 software package (Wavemetrics, Portland, OR, USA).^15,16^ Host (alanine, Ala) variant 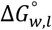 were previously determined and used to calculate 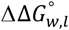 for each nonpolar guest variant.^15^ The *σ_NP_* for each site on OmpLA was determined by weighted linear regression using Igor Pro, with weights corresponding to the error (standard deviations) in 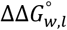:

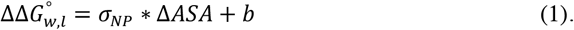

where

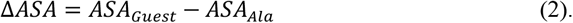

and *σ_NP_* and *b* are fitted parameters. The surface area for each nonpolar side chain was previously determined in the context of a Gly-X-Gly peptide.^16^

### Molecular Dynamics Simulations of OmpLA variants

All-atom molecular dynamics simulations of each OmpLA variant using the CHARMM36 force field were performed to determine the position of each guest side chain in the experimental DLPC bilayer. The monomeric crystal structure of OmpLA (1qd5) was used as the starting structure, and the remaining components of the system and scripts were created using CHARMM-GUI.^32–34^ A system was built for each OmpLA variant embedded in a DLPC bilayer and 0.2M NaCl at 37°C, resulting in 33 individual simulations. To approximate the experimental pH, some ionizable residues were protonated as described previously.^15^ Systems were initially equilibrated using the protocol provided by CHARMM-GUI, followed by an additional 50ns of equilibration to allow for the system to fully relax (Figure S1). All simulations used NAMD and were run on the Maryland Advanced Research Computing Center super computer.^35^

VMD plug-ins and homemade scripts were used to analyze all trajectories.^36^ To determine the position of the Cα for each variant relative to the phosphate plane, we first calculated the average *z*-position for the phosphate atoms in each leaflet of the bilayer. We then calculated the distance of the Cα for the variant side chain to the closest phosphate plane. These calculations were performed for each frame in the last 100ns of the trajectory, and the distances from the phosphate plane were binned into 0.1Å bins to create the histograms in Figure S2. Histograms were fit to Gaussian distributions using Igor Pro software to determine average positions and standard deviations found in Table S1.

To relate the Cα positions of each side chain to the local concentration of water ([water]) in the bilayer we derived an empirical relationship between [water] and *z*-position in the bilayer (Figure S3). These data were derived from simulations of a neat DLPC bilayer and extracted using the Density Profile Tool in VMD.^37^ We found that the data were well described by a sigmoidal function:

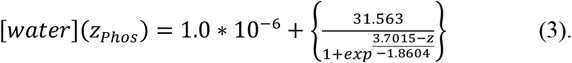

where *z* is the position relative to the phosphate plane of the bilayer.

The function can alternatively be adapted to any reference plane in the membrane by refitting [*water*] (*z*) with an adjusted z-position axis. For example, using the lipid carbonyl plane as the reference point (z = 0), the function changes to:

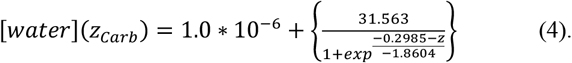

### Fitting and Derivation of *σ_NP_*([*water*|), [*water*] (*z*), *σ_NP_*(*z*), and 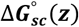

The relationship between *σ_NP_* and [water] was determined by weighted linear regression (errors derived from error of fitting *σ_NP_* using Equation 1) using Igor Pro, resulting in the following relationship:

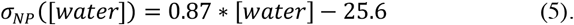

The [*water*] (*z*) (Equation 3) function allows for *σ_NP_* (*z*) to be derived:

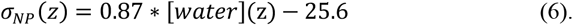

This function allows for *σ_NP_*(*z*) to be calculated at any *z*-position inside the phosphate plane of the bilayer. To derive 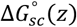 in kcal mol^-1^ for any nonpolar side chain from *σ_NP_*(*z*) the following relationship is used:

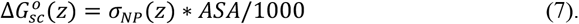

where ASA is the solvent accessible surface area for that side chain.

### Simulations and Analysis of Neat POPC Bilayer

A neat POPC bilayer (75 lipids per leaflet) was created using CHARMM-GUI with 0.2M NaCl at 37 °C. The CHARMM36m forcefield was used, and the simulation was equilibrated using CHARMM-GUI provided equilibration steps followed by 15 ns of additional equilibration time. The following 50 ns of simulation were used to measure the concentration of water molecules as a function of z-position in the bilayer and the z-position of the phosphate plane using the density profile plug in tool for VMD.^37^ The relationship between z-position and [water] was fit to a sigmoid function as described above. The *σ_NP_*(*z*) for POPC was also determined as described above.

## Results

### Experimental determination of a *z*-dependent nonpolar solvation function

In this work we define a function, *σ_NP_* (*z*), that relates the value of the *σ_NP_* to the *z*-position along the bilayer normal. This function is fundamentally derived from experimental protein folding titrations described below. Using a previously validated host-guest approach, we measured the thermodynamic stabilities of a series of host (Ala) and guest (nonpolar side chains isoleucine (Ile), methionine (Met), and valine (Val)) variants of the well-characterized membrane protein scaffold, E. *coli* outer membrane phospholipase A1 (OmpLA).^15,16,30,31^ Host/guest sites were chosen at varying positions across the bilayer to allow side chains to experience a range of local chemical compositions created by the amphipathic phospholipids in the membrane. In total, five lipid-facing sites located on well-defined transmembrane β-strands were used in this study (residues 120, 164, 210, 212, and 214) and are shown in Figure 1A. Figure 1B shows the water concentration gradient across the experimental DLPC bilayer and highlights the 60-fold change in water concentration accessed by these sites.

**Figure 1.**
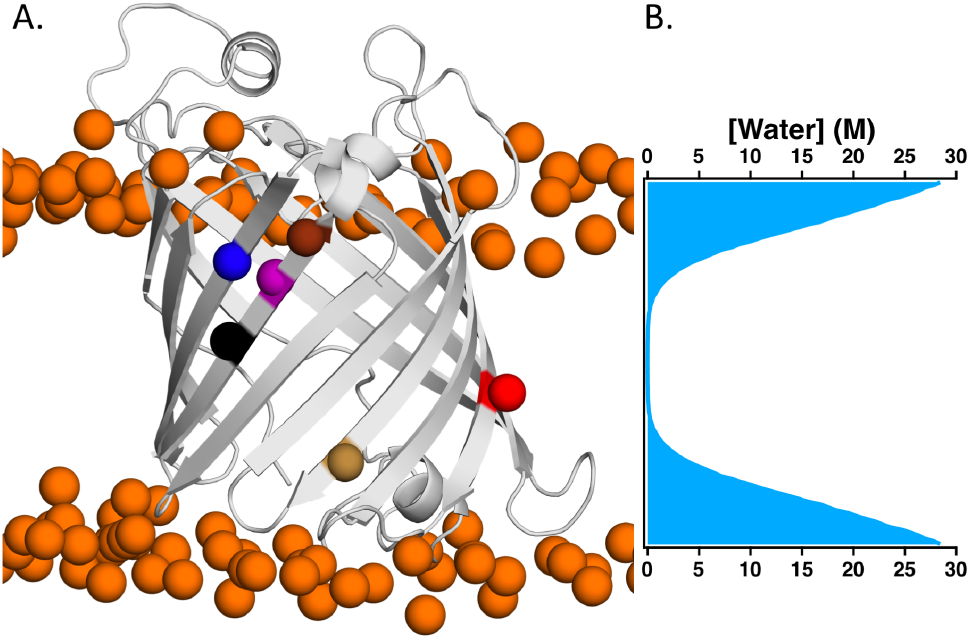
Host-sites on OmpLA exist in different water concentrations in the bilayer. **(A)** A snapshot of a molecular dynamics simulation of WT OmpLA in a DLPC bilayer is shown. Phosphate atoms of the DLPC bilayer are colored orange and six host sites on OmpLA used in this study are shown as colored spheres (black:210, yellow:164, red:120, purple:212, blue:223, brown:214). **(B)** The gradient of water concentrations inside the phosphate plane of the bilayer is plotted as a function of position in the bilayer. These values were obtained from previously published simulations of neat DLPC bilayers.^15^ This water gradient is aligned with the structure of OmpLA at the two phosphate planes. The sites on OmpLA were chosen because they sample a wide range of *z*-positions positions and chemical compositions.

The thermodynamic stabilities 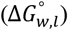 of all variants were measured using intrinsic tryptophan fluorescence-based chemical denaturation titrations (Figure S4).^15,16,18,31^ Figure S5 shows that each variant was enzymatically active, and Figure S6 shows that the folded and unfolded tryptophan fluorescence spectra for each variant overlay. Together these data indicate that single mutations do not affect the tertiary or quaternary structure of either the folded or unfolded states of OmpLA. We have previously shown that single site variants of OmpLA fold in a path-independent manner into large unilamellar DLPC vesicles, allowing for equilibrium thermodynamic parameters to be extracted by fitting the titration data to a three-state linear extrapolation model (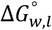 values listed in Table S2).^15,16,30^

The thermodynamic contributions of the guest side chain to folding were isolated by taking the difference in the 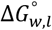 values of the host and guest variants at a given site on OmpLA as shown in the thermodynamic cycle in Figure S7. We interpret the resulting 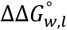 at each site to represent the water-to-bilayer transfer free energy change for each guest side chain relative to alanine at a particular site (Table S2). We find that our experimentally measured 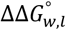 are remarkably similar to analogous, computationally estimated 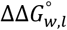 that were previously published by Liang and coworkers, though both Ile and Val deviated significantly at site 212 (Figure S8).^38^

### The nonpolar solvation energy is not constant along the bilayer interface

To determine the *σ_NP_* for each site on OmpLA we used the 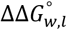 values for each nonpolar side chain. This dataset included previously published 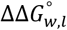 for leucine and phenylalanine.^15,16^ Figure 2 shows nonpolar 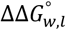 values plotted as a function of the nonpolar surface area of each side chain relative to alanine (Δ*ASA*) for each site. The *σ_NP_* at each site on OmpLA is the slope of the linear fit of the correlation between nonpolar 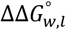 and Δ*ASA* (Equations 1 and 2, fit parameters in Table S3). The *σ_NP_* values vary nearly two-fold from site to site with a range that spans from −27.3 to −16.1 cal mol^−1^ Å^−2^.

**Figure 2.**
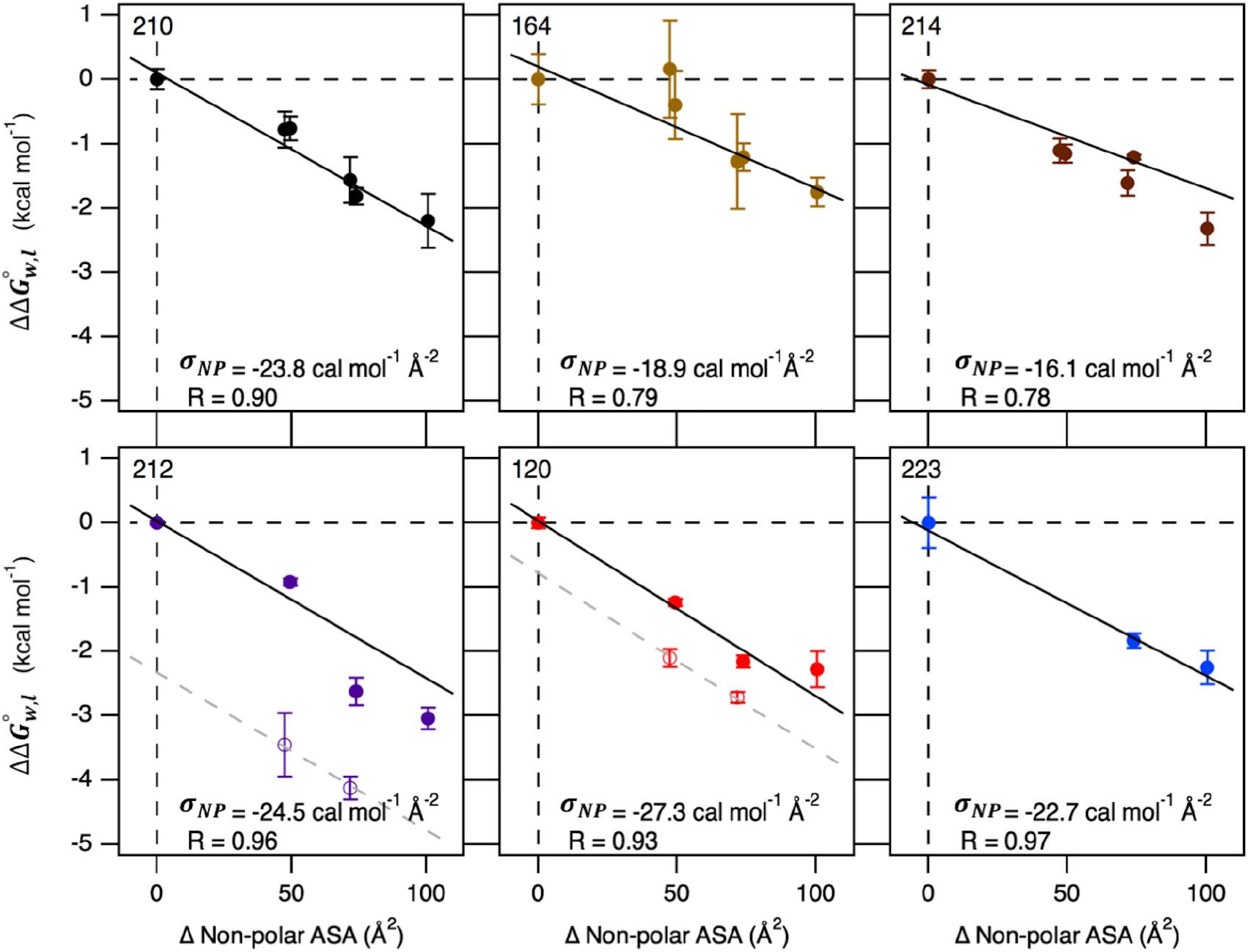
Nonpolar solvation parameter is not constant throughout the bilayer. The nonpolar solvation parameter, *σ_NP_*, was determined for each site on OmpLA by calculating the slope of the linear relationship between 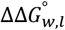 for each nonpolar side chain at that site with the change in buried surface area compared to alanine in a Gly-X-Gly peptide (Equation 1).^16^ Linear fits were weighted by the error in 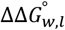 (standard deviations found in Table S2) for each site, with the slope of the line (*σ_NP_*) and the Pearson correlation coefficient reported for each fit shown at the bottom of each panel. For sites 212 and 120, beta-branched side chains isoleucine and valine were omitted from *σ_NP_* determination as they had anomalously greater 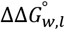 compared to the other nonpolar side chains. The *σ_NP_* determined from non-beta-branched side chains (solid line in 212 and 120 panels) fits the beta-branched side chains (dotted line), indicating that the increase in energy is derived from favorable local interactions that are restricted to beta-branched residues. By taking the difference in the y-intercept for the two lines, we estimate the energy gained due to local interactions for beta-branched residues at these sites to be 2.38 kcal mol^−1^ for site 212 and 0.80 kcal mol^−1^ for site 120. The *σ_NP_* for site 223 was determined only using previously collected data.^16^

This experimental setup requires a reference host side chain whose energetic contributions must be removed to obtain a reference-free scale. As alanine is the amino acid side chain in the host protein, we obtain a reference-free scale by removing the intrinsic contributions of Ala to 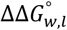 at each site using the site-specific *σ_NP_* values. The reference-free 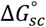 for alanine at each host site was determined by multiplying the corresponding *σ_NP_* by the ASA of the alanine side chain.^16^ The reference-free 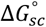 for the other nonpolar side chains were calculated by adding the 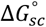 for alanine to the 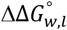 at each site. The 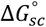 for each nonpolar side chain, including alanine, are listed in Table S4.

The nonpolar solvation parameter is linearly correlated with the amount of water in the bilayer. We next addressed the question of how the local chemical composition of the bilayer modulates the *σ_NP_* by deriving the relationship between the *σ_NP_* and *z*-position along the bilayer normal. This results in the function *σ_NP_*(*z*). To accomplish this, we calculated the *z*-position of the side chain Cα of each OmpLA variant from all-atom molecular dynamics simulations conducted in the experimental lipid bilayer. Previous work showed that introduction of single side chains has no discernable effect on the *z*-position or tilt of OmpLA in the bilayer.^29^ Figure S1 shows that each simulation was equilibrated after 50ns, and analyses were performed on the final 100 ns of each run. Instead of using the bilayer center as the reference point, (*z* = 0), we calculated the position of each Cα relative to the average position of the phosphate plane of the bilayer.

We chose the phosphate plane as our reference point for z because it can be more easily applied to bilayers with different lipid compositions and acyl chain lengths. The carbonyl plane is an alternative reference coordinate employed in the literature and could similarly be used. Figure S2 shows that variant Cα positions are generally well described by Gaussian distributions and that our sites capture the majority of the region between the bilayer center and phosphate plane. Table S1 lists the mean *z*-position and standard deviations for each variant Cα, which were averaged to calculate the *z*-position for the σ_NP_ at each site. We found that the bilayer *z*-position dependence of water concentration in our experimental DLPC bilayer was described by a sigmoidal function (Equation 3, Figure S3).^15^ Using this correlation we calculates the corresponding local water concentration for each *σ_NP_* (Table S5).

Figure 3A shows that the *σ_NP_* is linearly correlated with the water concentration in the bilayer (Equation 5, R^2^ = 0.84). The ~5,000-fold change in water concentration illustrates the large range of solvent conditions accessed in our experiments and necessitates presentation using a logarithmic abscissa. This novel functional relationship we derived from our measurements using a native protein and a phospholipid bilayer confirms our hypothesis that the water in a bilayer is a reporter on the hydrophobicity of the membrane and that the local hydrophobicity modulates membrane protein stabilities through the *σ_NP_* directly.

**Figure 3.**
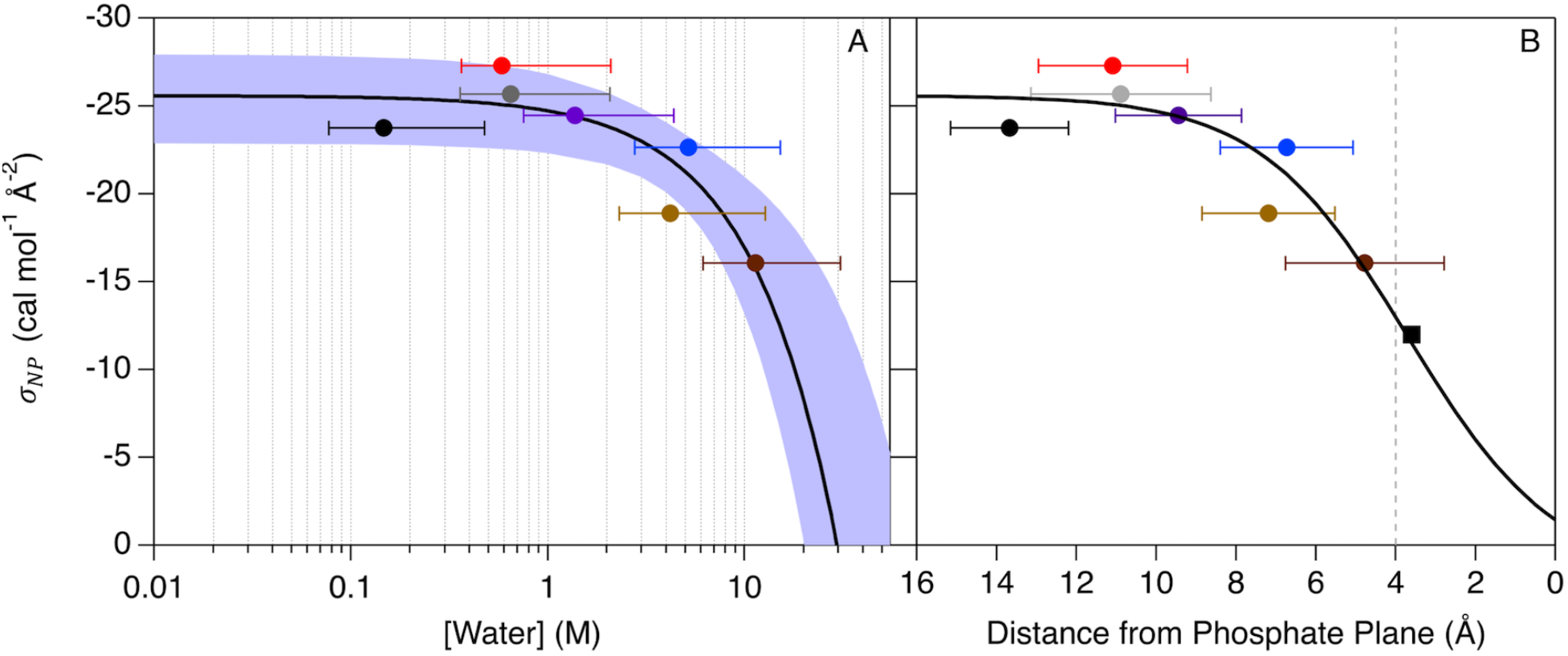
Nonpolar solvation parameter is linearly correlated with water concentration in the bilayer. The water-to-bilayer *σ_NP_* calculated at each host site in OmpLA (colored as in Figure 1) and PagP (gray point) plotted as a function of water concentration (Table S4) (**A**) and distance from the phosphate plane (**B**).^18^ The *σ_NP_* are colored according to the scheme in Figure 1, with the *σ_NP_* derived from PagP position 111 colored gray. The position of *σ_NP_* for a given site is the average of the positions of each side chain at that site and the error bars reflect the standard deviation (Table S1, NP row). *σ_NP_* and water concentration are linearly correlated with the equation shown in the bottom right hand corner of Panel **A** (black line: R^2^ = 0.84; 95% confidence interval shaded in light blue). Using the derived relationship between bilayer position and water concentration shown in Figure S3, the direct relationship between *σ_NP_* and bilayer position can be determined (Panel **B**). We were also able to assign a previously measured “water-to-interface” solvation parameter (−12 cal mol^−1^ Å^−2^) to an exact position in the bilayer (3.75 Å from the phosphate plane) using this function (shown as a black square in Panel **B**) and find that it is reporting on the position of the lipid carbonyl plane in the bilayer (dashed vertical gray line).^17^ The position-dependent function describing *σ_NP_* allows for the 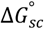 of any nonpolar side chain to be determined for any position of the bilayer inside the phosphate plane (shown in Figure 4).

It is also instructive to consider the *σ_NP_* using bilayer *z*-coordinates. Figure 3B shows the bilayer z-position dependence of *σ_NP_* in which the line of best fit in Figure 3A is transferred to units of distance from the phosphate plane of the membrane. Using the resulting function, *σ_NP_*(*z*) (Equation 6), the *σ_NP_* can easily and accurately be determined at any position in the bilayer. *σ_NP_*(*z*) can also be applied to any membrane system with a known [*water*] gradient along the bilayer normal. Figure S9 shows the [water](z) and *σ_NP_*(*z*) functions for a neat POPC bilayer, which is almost indistinguishable from DLPC indicating *σ_NP_*(*z*).

The *σ_NP_*(*z*) function plateaus to a value of −25.6 cal mol^−1^ Å^2^ in the anhydrous core of the bilayer, which agrees with the *σ_NP_* determined for this region of the bilayer in previous studies.^13,16,18,19^ We find that the magnitude of the *σ_NP_*(*z*) decreases in the membrane interface to a predicted *σ_NP_* equal to −1.4 cal mol^−1^ Å^−2^ at the phosphate plane, which is much less favorable than previous estimates of σ_NP_ for the interface (−12 cal mol^−1^ Å^−2^).^17^ Using *σ_NP_*(*z*), we calculate that this previously measured *σ_NP_* for the water-to-interface transition corresponds to a z-position approximately 3.75 Å inside the phosphate plane. This *z*-position corresponds to the approximate position of the lipid carbonyl plane, which is a commonly used reference as it is approximately the boundary between hydrocarbon and interfacial regions of the bilayer.

### Bilayer *z*-position dependence of reference-free 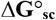

The large change in the *σ_NP_* across the bilayer interface significantly impacts the contribution of nonpolar side chains to the overall stability of membrane proteins in a position-dependent manner. This can be visualized by using the *σ_NP_*(*z*) to calculate a reference-free, nonpolar 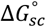 as a function of *z*-position in the bilayer (Equation 7). Figure 4 shows the 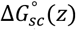 for each nonpolar side chain (black lines) and the 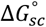 for each site on OmpLA determined above (points). Side chains located in the bilayer interface can be several kcal mol^−1^ less favorable than side chains in the dehydrated core of the bilayer. Additionally, our findings indicate that interfacial side chains separated by a single turn of an alpha helix can have dramatically different contributions to the overall stability of a membrane protein.

**Figure 4.**
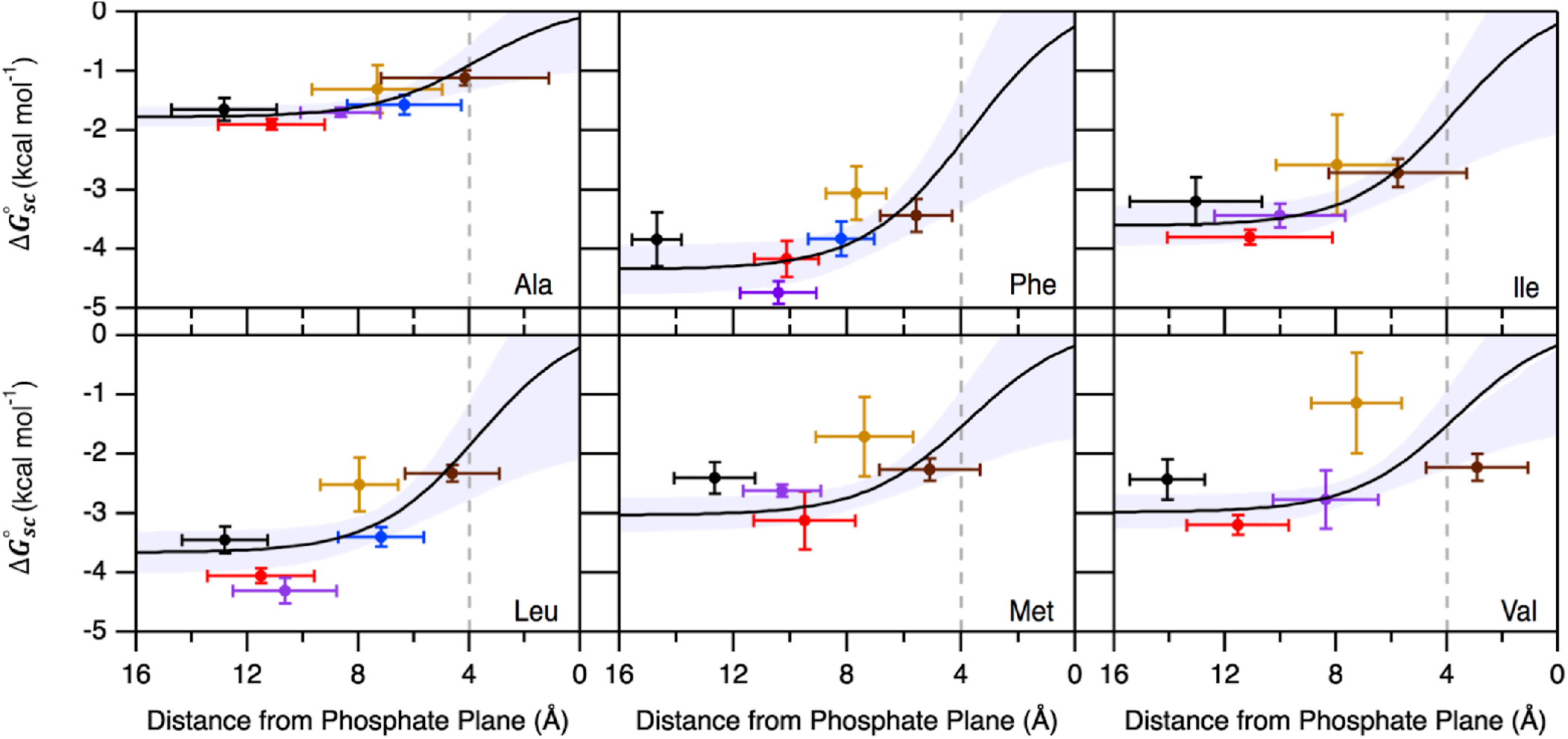
Bilayer position dependence of nonpolar 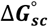. Reference-free side chain transfer free energies are plotted as a function of the average distance from the bilayer phosphate plane, as determined from molecular dynamics trajectories. 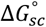 for each host site in OmpLA are colored as in Figure 1. Error bars for both 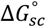 and bilayer position represent the standard deviations. The solid black line in each panel represents the simulated 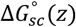 profile for each nonpolar side chain. This function is derived from *σ_NP_*(*z*) multiplied by the nonpolar surface area of each side chain (Equation 7). Vertical dotted lines represent the position of the lipid carbonyl groups.

The functional free energy dependence for the alanine side chain, 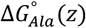, is particularly useful because the experimental setup necessitates that all side chain free energy perturbations be measured relative to Ala. Upon subtraction of the reference (Ala) energy, the *z*-position-dependence for any side chain can be place on a reference-free scale. Accordingly, the data herein allows conversion of previously measured 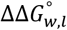 values for arginine, tryptophan, and tyrosine to reference-free 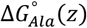 scales.^15,16^ Following the protocol above, a combination of new (Arg) and published (Tyr and Trp) allatom molecular dynamics simulations were first used to determine the *z*-position of the Cα atom for each variant.^15^ This 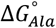 value at a particular *z*-position was calculated using the alanine functional free energy dependence 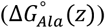, and this value was added to the corresponding experimental value for the guest side chain.

Figures S9, S10 and Table S6 shows these adjustments applied to the arginine, tryptophan, and tyrosine side chains.^15,16^ As expected, these data show that there is a larger energetic penalty for the placement of the arginine Cα within the bilayer ~13 Å from the phosphate plane (+2.5 kcal mol^−1^). This observation is in contrast to a small, favorable energetic contribution when the side chain is located in the bilayer interfacial region (−0.5 kcal mol^−1^). The 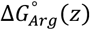 is remarkably similar to the reported translocon-mediated profile from Arg insertion in a α-helical TMD.^14^ This similarity most like is derived from the fact that Arg can snorkel to the interface at any position in a TMD, which would change its thermodynamic “end state” to be the interface in all cases. The energetic trends for the aromatic side chains 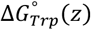 and 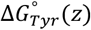 reveal an opposite but also expected behavior: both side chains show an energetic preference for the bilayer interface as compared to the nonpolar central region of the bilayer.

### The translocon energetically mimics the bilayer interface to facilitate TMD insertion into the membrane

With the reference-free, water-to-bilayer 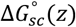 for nine amino acids determined as a function of *z*-position in the bilayer, we next asked whether our findings could be used to better understand membrane protein folding by the Sec translocon machinery. The translocon acts as a protein channel in the bilayer that allows α-helical TMDs to partition into the bilayer while protecting hydrophilic loops and turns from exposure to the hydrophobic membrane.^10,39^ All-atom molecular dynamics simulations have identified that the chemical properties of water in the pore of the translocon mimic the properties of water in the interfacial region of the membrane.^40^

Thus, a hypothesis for translocon function is that it mimics the thermodynamics of an interface-to-bilayer transition, not a water-to-bilayer transition.^10^ Figure 5A shows the relevant free energy reactions that can be written for these processes. The water-to-bilayer-center nonpolar 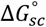 values reported here are much greater in magnitude than the Δ*G_app (t,b)_* for the translocon-to-bilayer transition.^14,41^ Because we have measured the water-to-bilayer 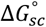 across the bilayer normal, we can calculate the interface-to-center transition by taking the difference between 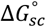 at the center of the bilayer (A210) and the 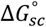 at the most interfacial site (Y214) (Figure 5A). Figure 5B shows that our interface-to-bilayer 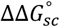, or 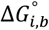, agree remarkably well with the Δ*G_app (t,b)_*.^41^ Additionally, the *σ_NP_* for the interface-to-center transition (*σ_NP,int–cen_* = *σ_NP,214_* – *σ_NP,210_*) is equivalent to the σ_NP_ calculated for translocon-mediated folding (7.7 and 6-10 cal mol^−1^ Å^2^, respectively).^26^ The remarkable energetic equivalence between the interface-to-center and translocon-mediated transfer free energies provides thermodynamic backing to the hypothesis that the translocon aids TMD folding by mimicking properties of the interface.

**Figure 5.**
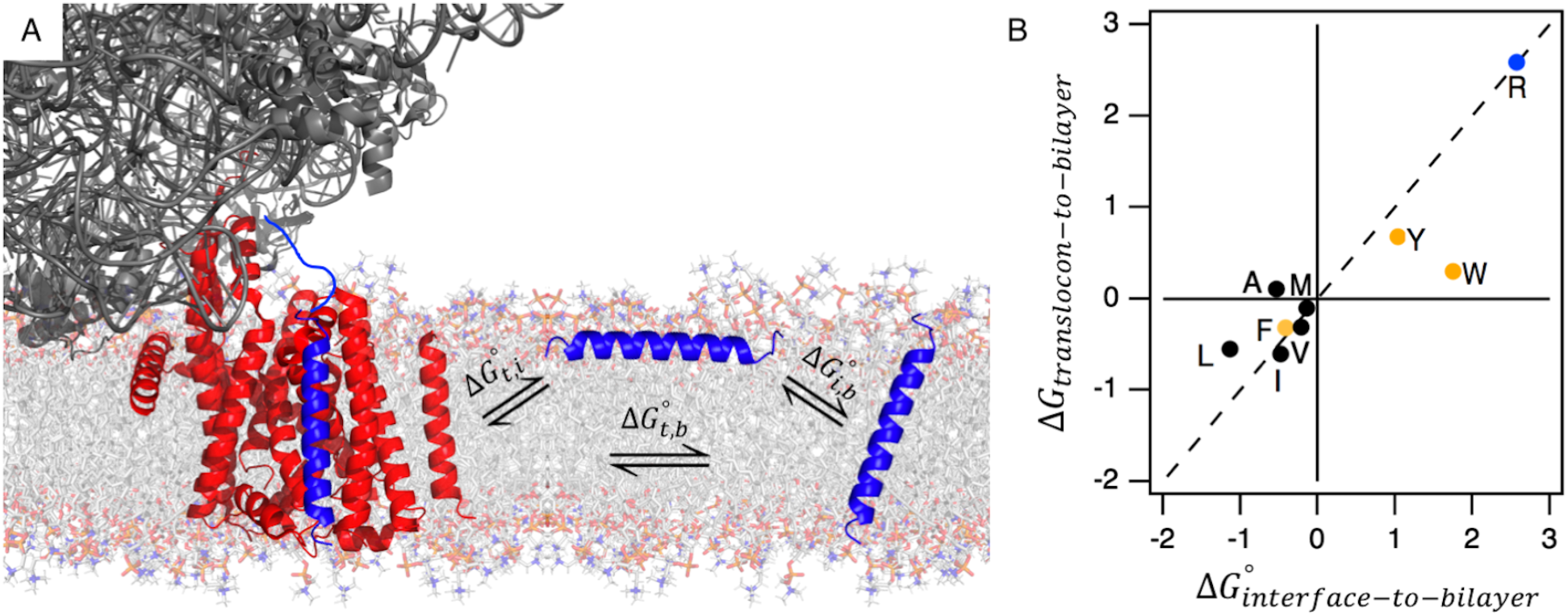
The translocon energetically mimics the bilayer interface. **(A)** A cartoon schematic of the co-translational insertion of a helix (blue) via the translocon (ribosome colored dark gray, translocon colored red; PDB code: 5GAE) is shown on the left. The helix can either partition to an interfacial conformation, which is energetically described by 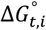, or to a transmembrane conformation, described by 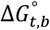 (*t* = translocon, *i* = interfacial, *b* = transmembrane). The interfacial-to-transmembrane conformation, which can occur in the absence of the translocon, is energetically described by 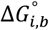. **(B)** The values for 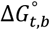 and 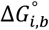 are plotted for nine side chains, with the dotted line representing energetic equivalence between the two equilibria (black = nonpolar, gold = aromatic, blue = ionizable).^41^ For all side chains, except Trp, 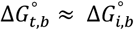, indicating that the translocon energetically mimics the interface 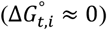. The deviation for Trp is hypothesized to be due to the energetic preference of Trp to exist in the bilayer interface^15,42^.

## Discussion

The thermodynamic stability of a membrane protein regulates its structure, conformational landscape, and ability to perform its biological function.^7,11^ A key determinant of this stability is the 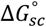 of each side chain from water into the membrane. Historically, experiments measuring side chain transfer from water to organic solvents such as cyclohexane or octanol were used to approximate 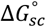.^19,22^ While these experiments accurately describe the water-to-bilayer-center transition, they failed to capture thermodynamic properties resulting from the dramatic chemical heterogeneity of the interfacial region of the bilayer.

In this study we have measured the 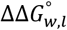, for Ile, Met, and Val as a function of z-position in the bilayer. Using these 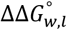, along with previously measured 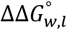, for Leu and Phe we were able to determine a novel functional form for the *σ_NP_* across the bilayer normal, *σ_NP_*(*z*).^15,16^ We discovered a direct correlation between the *σ_NP_* - and thus the 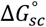 for nonpolar side chains - with the local water concentration in the bilayer. The *σ_NP_* (*z*) is a convenient numerical expression for assessing the magnitude of the hydrophobic effect, as it relates nonpolar surface area to the energy gained from transferring the surface area from bulk water to a dehydrated environment.

Importantly, the dependence of nonpolar 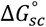 on local water concentration differs starkly from the water-dependence of backbone hydrogen bond (bbHB) formation in membrane proteins. Although it was long expected that bbHB energies would also be strongly influenced by the steep water gradient intrinsic to bilayer interface, we recently observed bbHB energies to be constant throughout this region.^42^

The experimental data here offer an opportunity to consider how the translocon insertion process relates to the underlying physical chemistry of the reaction. Our data provides evidence that translocon-mediated membrane protein folding energetically mimics the interface-to-bilayer transition for 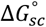. We find that 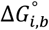 is approximately equal to Δ*G_app (t,b)_* for eight of the nine side chains investigated, with Trp being the only outlier; we speculate that this one outlier is most likely due to differential aromatic interactions in the OmpLA and translocon experimental setups.^15,41^ This finding provides thermodynamic support for a recent model for translocon function in which transmembrane α-helices sample both interfacial and translocon associated conformations before they insert into the membrane.^10^ Our findings indicate that these two states would be essentially energetically equivalent, indicating the energy barrier between these two conformations may be nonexistent with switching between the two states determined by the kinetics for a given TMD sequence.

Our results should improve the speed and accuracy of membrane protein structure prediction and design algorithms.^43–45^ As membrane proteins account for over half of all therapeutic drug targets,^46^ having accurate force fields to describe membrane protein energetics is essential. This simple, linear function *σ_NP_*(*z*) could be adapted to implicit membrane models or could alternatively be applied explicitly in systems such that a local value of *σ_NP_* to be calculated over water concentrations ranging from millimolar to 30 molar. Subsequently, better estimates of the energetic contribution of any lipid-facing, nonpolar side chain to the stability of membrane protein.

By relating the *σ_NP_* to the chemical properties of the bilayer, the *σ_NP_*(*z*) can be customized to any bilayer lipid composition by calculating the water concentration gradient of that particular bilayer (illustrated in Figure S9 for a POPC bilayer). We find that the *σ_NP_*(*z*) for both POPC and DLPC bilayers are almost indistinguishable, indicating that lipid acyl tail length and saturation have little effect on the shape and magnitude of *σ_NP_*(*z*) across the bilayer interface. Further work is needed to assess the impact of headgroups and other macromolecules found in bilayers such as cholesterol on the shape of *σ_NP_*(*z*). In addition, one caveat of *σ_NP_*(*z*) is that it relies on additive forcefields, which may underestimate the local water concentration in the bilayer interface. As polarizable forcefields become more readily accessible the *σ_NP_*(*z*) could be refined to even more accurately quantify the energetics of nonpolar side chain burial in membranes.

The *σ_NP_*(*z*) and the nine 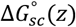 functions derived here also could be applied to predicting the contribution of the binding energy of proteins to the bilayer interface (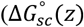 functions found in Table S7). By relating energy to *z*-position of side chains in the bilayer, binding energies can be tuned based on structural information derived from experiment or computation. The simplest application of the 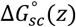 functions would apply to proteins that do not undergo large structural changes upon membrane interaction, as the binding energy would be dominated by lipid-side chain interactions. In cases of conformational changes, our values may find utility in providing baseline values for the partitioning of many membrane binding proteins (such as antimicrobial peptides) even when the global binding energy contains additional contributions from other sources, such as the energy of the coil-to-helix transition that occurs upon binding.^16^ Given the complexity of membrane protein folding, even for apparently simple peptides, the safest application of our values is in understanding the effect of a point mutation on bilayer-protein interactions.

Additionally, our findings delineate how the different transfer free energies reported here should be applied to various biological functions. The critical parameter for consideration is the endpoints for the reaction at hand. The Δ*G_app (t,b)_* or 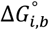 can be used to approximate the contribution of a side chain to the folding of alpha-helical TMDs.^14,41^ On the other hand, water-to-bilayer 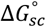 are more applicable to modeling both changes in stability of different conformers of a membrane protein, the transition between interfacial and transmembrane conformations of antimicrobial peptides, and potentially unfolding processes, such as extraction of transmembrane regions from the bilayer by AAA-ATPases.^47^ Experimentally determining these thermodynamic parameters for membrane proteins is extremely challenging and accurate computational modeling will allow for greater understand of essential processes such as the gating of a channel or the cycling of a transporter.^48–50^ At the very least, we anticipate that 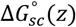 can be used to better understand the effects of mutations that may impair membrane protein folding such as those occurring in cystic fibrosis and Charcot-Marie-Tooth disease.^51^

### Conclusion

In summary, we have determined the relationship between nonpolar side chain transfer free energies, solvent accessible surface area, and the local water concentration in the bilayer. Using this relationship, we calculate 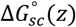 functions for nonpolar side chains and remove the alanine dependence of previously measured 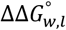 for Arg, Trp, and Tyr. Our work shows that the translocon energetically mimics the bilayer interface for side chain transfer into the membrane. Together, these findings increase our understanding of the driving forces of protein stability in membranes by essentially quantifying the hydrophobic effect along the bilayer normal and should increase the accuracy of computational workflows to identify, design, and understand the energy landscapes of membrane proteins.

## ASSOCIATED CONTENT

### Supporting Information

The Supporting Information is available free of charge on the ACS Publications website.

The Supporting Information contains representative chemical denaturation titrations, spectroscopic and enzymatic confirmation of OmpLA variant folding, thermodynamic cycle, molecular dynamics simulation analysis, reference-free transfer free energy profiles for Arg, Trp, and Tyr, stabilities of OmpLA variants, side chain transfer free energy values (PDF).

## Author Contributions

D.C.M. and K.G.F. designed research, D.C.M. performed experiments and analyzed data, D.C.M. and K.G.F. wrote the paper. All authors have given approval to the final version of the manuscript.

## Notes

The authors declare no competing financial interest.

## ACKNOWLEDGMENTS

The authors would like to thank Dr. Sarah McDonald and Margo Goodall for assistance with cloning the variants used in this study, Mark Kreutzberger for growth and purification of some OmpLA variants used in this study, and Dr. Sarah McDonald and Dr. Patrick Fleming for experimental and computational guidance.

## ABBREVIATIONS

OmpLA: E. *coli* outer membrane protein phosphatase A1;
PagP: E. *coli* lipid A palymitoyltransferase PagP;
DLPC: 1,2 didodecanoyl-sn-glycero-3-phosphocholine;
LUV: large unilamellar vesicle;
TMD: transmembrane domain;
σ_NP_: nonpolar solvation parameter;
Δ*G*^∘^_*w,l*_: OmpLA variant free energy of folding;
ΔΔ*G*^∘^_*w,l*_: alanine-referenced, water-to-bilayer side chain transfer free energy;
Δ*G*^∘^_*sc*_: reference-free, water-to-bilayer side chain transfer free energy;
Δ*G_app (t,b)_*: translocon-to-bilayer side chain transfer free energy;
Δ*G*^∘^_*i,b*_: interface-to-bilayer side chain transfer free energy;
Δ*G_t,i_*: translocon-to-interface side chain transfer free energy.

---

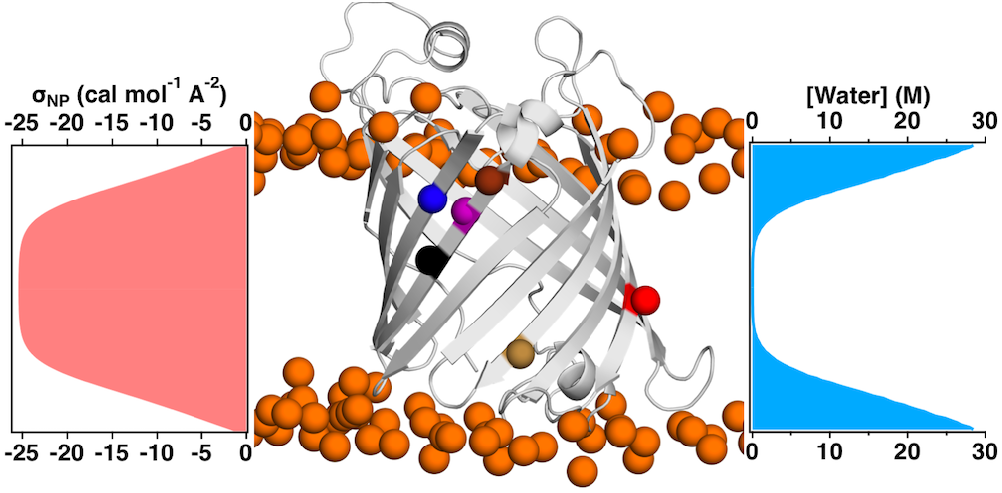

