## Supplemental Information for "Local Bilayer Hydrophobicity Modulates Membrane Protein Stability"

Dagan C. Marx and Karen G. Fleming*
Thomas C. Jenkins Department of Biophysics, Johns Hopkins University, Baltimore, MD.

Corresponding Author: Karen G. Fleming


### Supplemental Figures

*Note –Supplemental Figures were created using Igor Pro (WaveMetrics, Lake Oswego, OR, USA)*

**Figure S1. Molecular dynamics simulations of OmpLA variants are equilibrated after 50 ns**

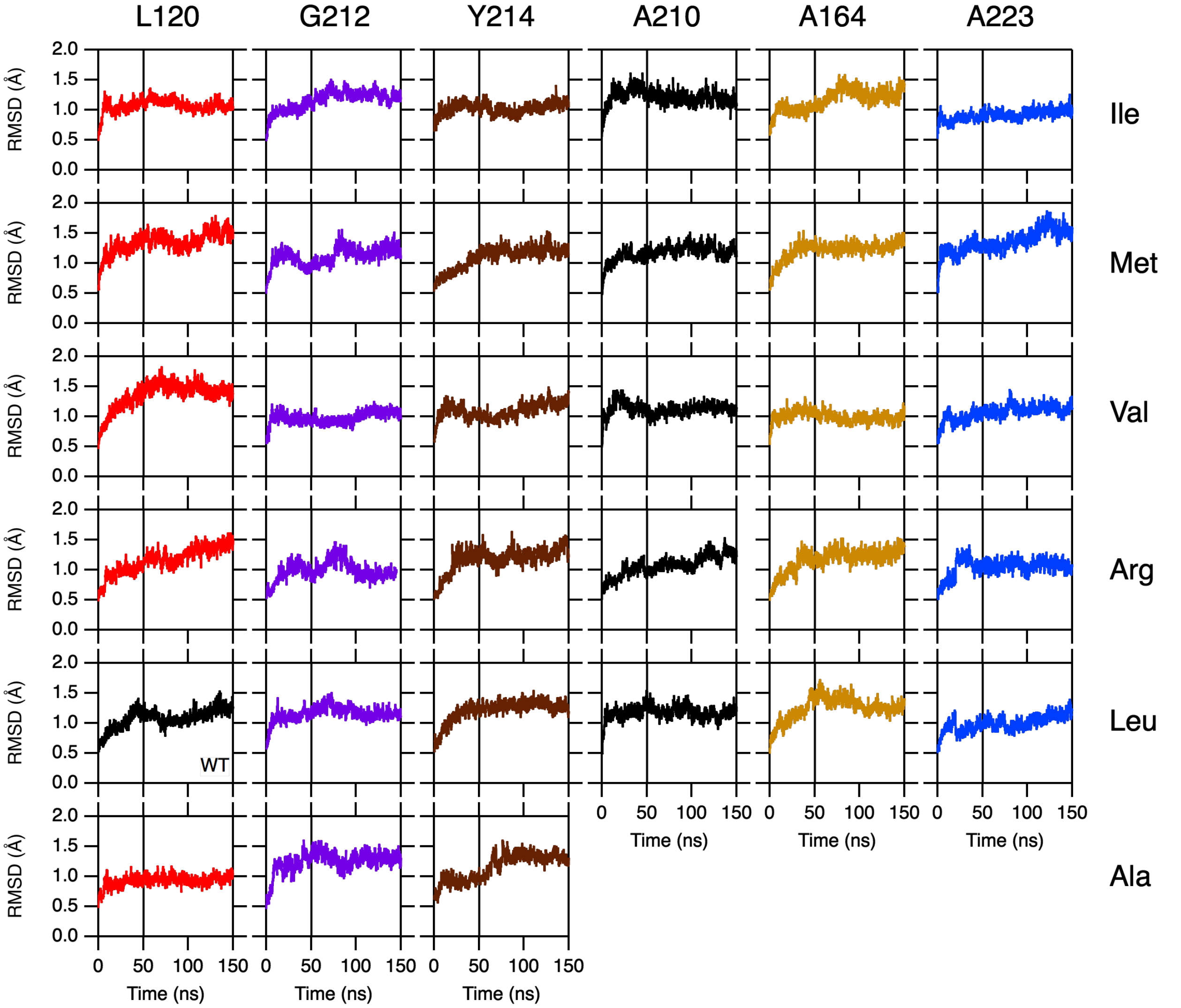

This figure shows the equilibration of each 150 ns molecular dynamics trajectory by calculating the RMSD for the transmembrane beta-sheet of OmpLA as a function of time. RMSD plots are colored as in Figure 1. Further analysis of the trajectories was performed using the last 100ns of each trajectory.

**Figure S2. Histograms of side chain Cα position relative to phosphate plane for each OmpLA variant**

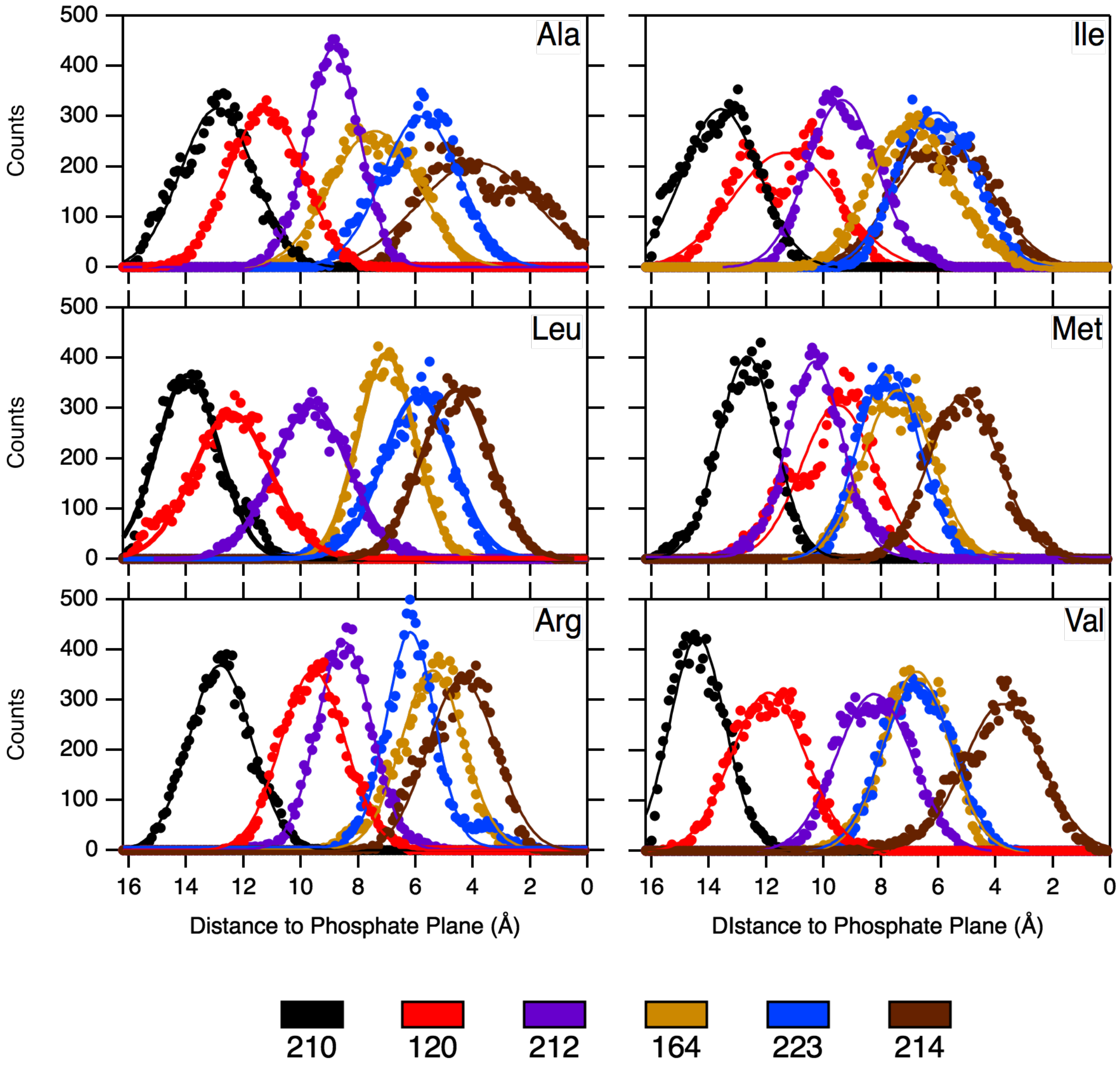

Cα positions for each variant were calculated for every time-step and binned by a 0.1Å step size over the last 100ns for each trajectory. Histograms relating the position of each variant in the bilayer relative to the phosphate plane are shown above. Solid lines are fits to a Gaussian distribution, from which the average position and error in the form of a standard deviation were extracted (Table S3). Site 223 overlaps in position with sites 164 and 214, which is why $\Delta\Delta G_{w,l}$ for Ile, Met, and Val were not measured at this site.

**Figure S3. The relationship [water] and bilayer z-position is well described by a sigmoid function**

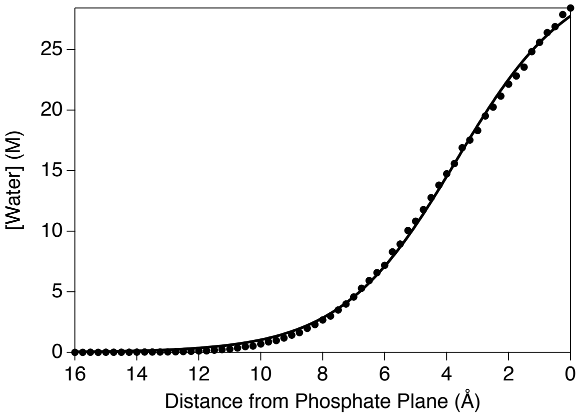

The data shown here (black circles) are derived from previously published molecular dynamics simulations of a neat DLPC bilayer using the density profile plugin.[1] The data are well described by a single sigmoid indicated by the solid line. (Equation 4).

**Figure S4. Representative Folding Titrations for Ile, Met, Val OmpLA Variants**

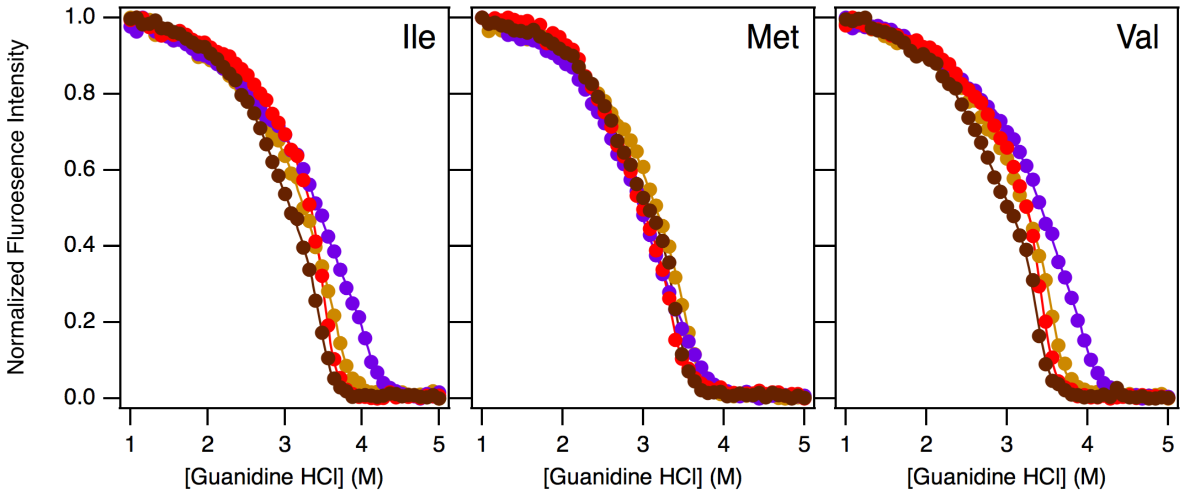

Representative chemical denaturation titrations are shown for each hydrophobic variant of OmpLA. Sites are colored as in Figure 1 (L120:red, A164:tan, G212:purple, Y214:brown). Fits to a three-state linear extrapolation model are shown as lines in the same color as the data set to which they are fit.[2–4]

**Figure S5. Nonpolar Variants of OmpLA are Enzymatically Active**

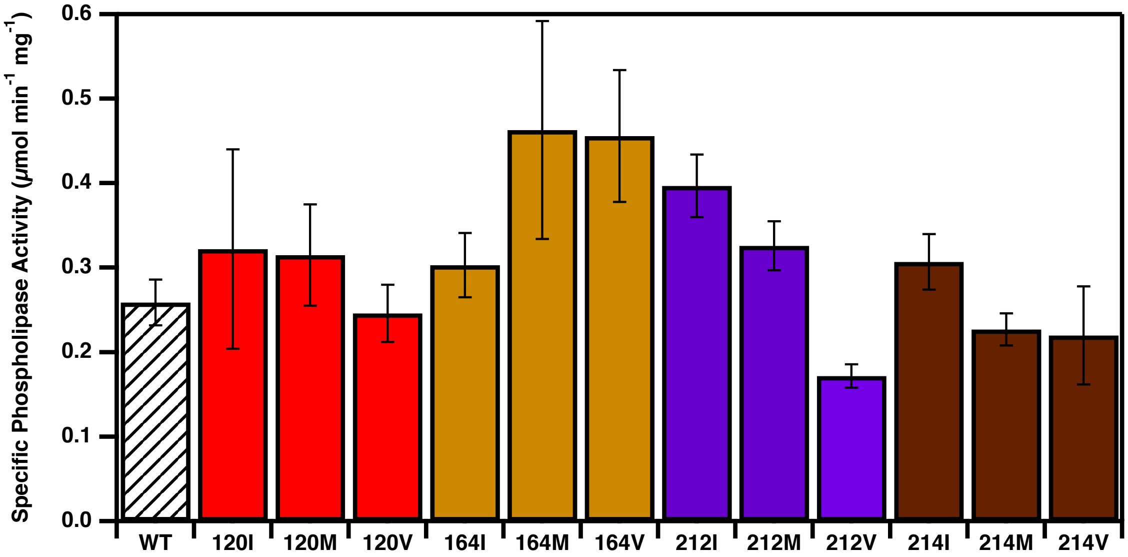

The activity of each new OmpLA variant was determined using a phospholipase activity assay on protein folded into DLPC LUVs in 1M Guanidine HCl.[2,3] The specific activities of each nonpolar guest-variant used in this study, and WT OmpLA, are shown colored as in Figure 1. Each variant has measurable enzymatic activity similar to WT, indicating that each variant is able to fold to a WT-like structure in DLPC bilayers. Error bars represent standard deviations from three independent trials.

### Figure S6. Nonpolar Variants of OmpLA Fold into DLPC LUVs

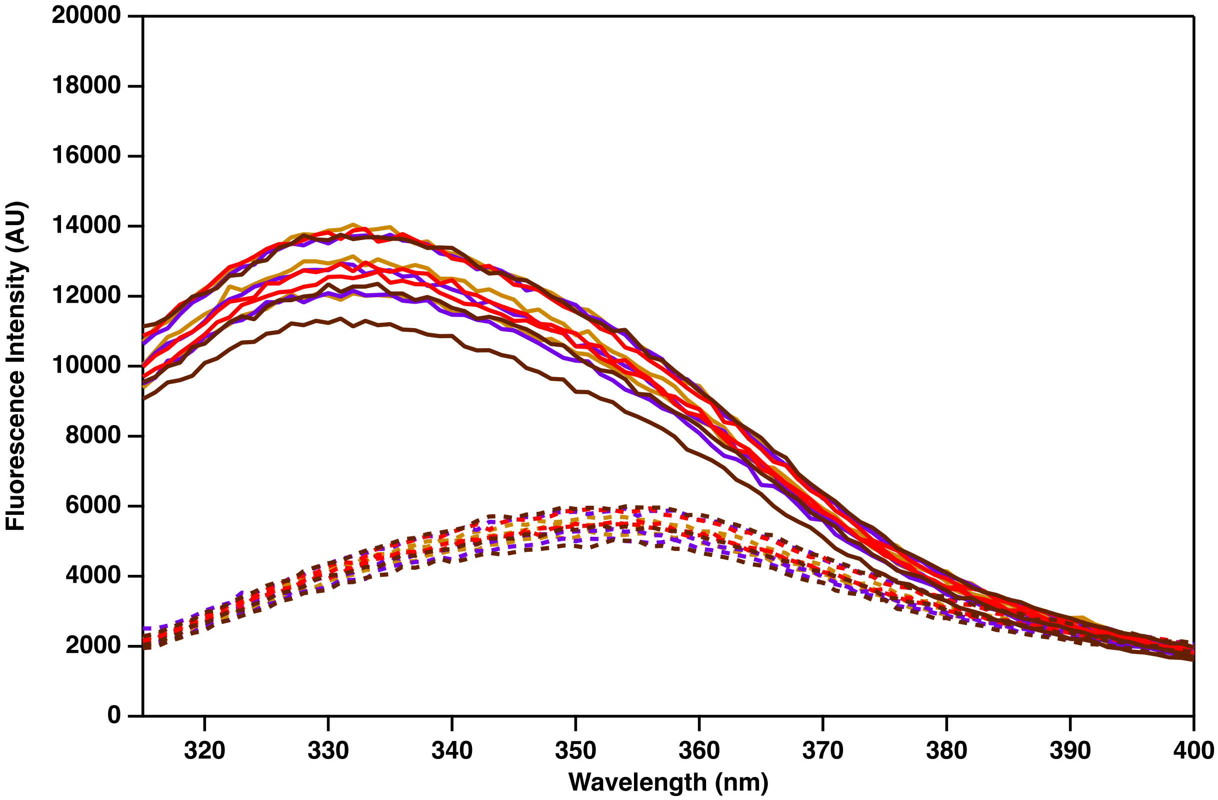

Tryptophan-fluorescence wavelength scans for OmpLA variants used in this study are shown. Solid lines correspond to folding conditions (1M GdnHCl) and dotted lines correspond to denaturing conditions (5M GdnHCl). The similarity of the shape and signal intensity for each OmpLA variant indicates that they all adopt similar structures in both folded and denatured conformations.

**Figure S7. Schematic of the host-guest calculation of $\Delta\Delta G°_{w,l}$.**

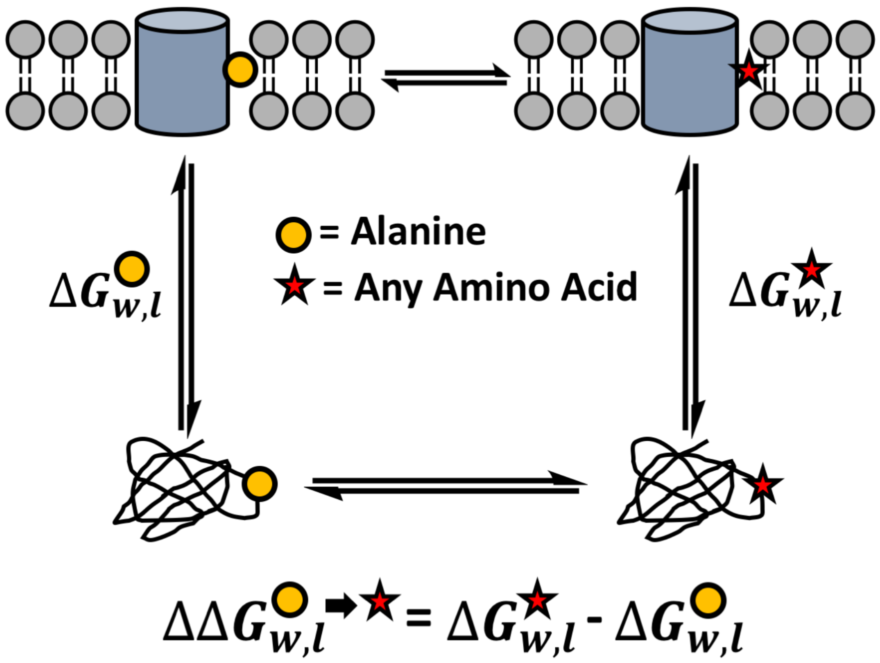

The stability ($\Delta G^{o}_{w,l}$) of both host (alanine) and guest (any other amino acid) variants at a site of OmpLA are measured using chemical denaturation titrations. The guest side chain transfer free energy with respect to alanine, $\Delta\Delta G^{o}_{w,l}$, is calculated by taking the difference in the host and guest $\Delta G^{o}_{w,l}$.

**Figure S8. Comparison of Experimental and Computational $\Delta\Delta G^{o}_{w,l}$ for Ile, Met, and Val.**

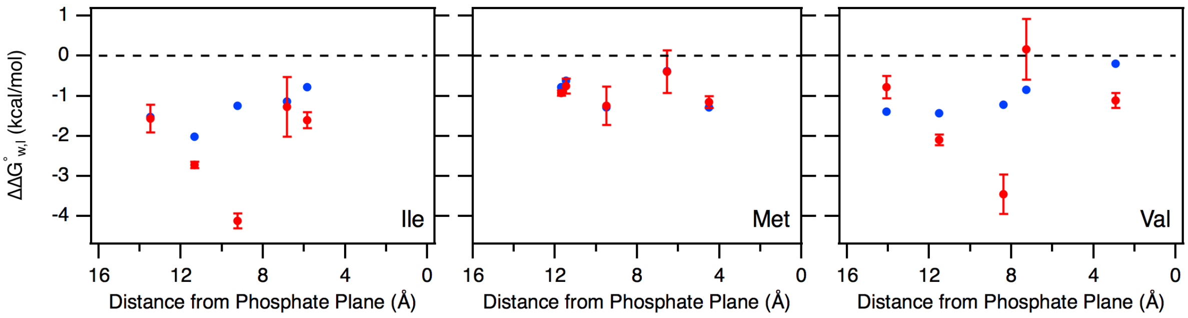

Comparison of experimentally measured $\Delta\Delta G^{o}_{w,l}$ for Ile, Met, and Val (red points) with computationally estimated $\Delta\Delta G^{o}_{w,l}$ (blue points) at these positions.[5] Error bars for experimental data points correspond to standard deviations (Table S1).

**Figure S9. Flowchart for Applying $\sigma_{NP}(z)$ to Other Phospholipid Bilayers (Example of POPC)**

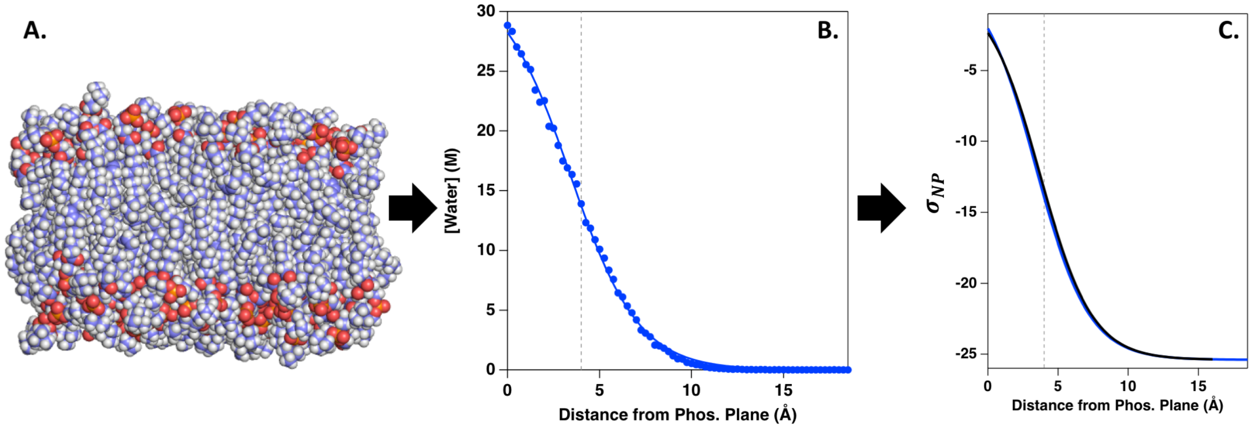

This flowchart details how the relationship between the $\sigma_{NP}$ and either water concentration or bilayer z-position can be applied to other bilayers using a POPC bilayer as an example. **(A)** First, the bilayer must be constructed properly and equilibrated using molecular dynamics simulations to ensure that the chemical composition of the bilayer interface is as accurate as possible. **(B)** Bilayer properties can be calculated using the density plug-in tool in VMD.[1] To calculate the water density distribution, we calculate the average density of the oxygen atoms in the water molecules as a function of z-position using all frames from the 50ns simulation (blue circles). The relationship between [water] and z-position ($[water](z)$) can be fit to a sigmoid function (Equation 3; blue line). **(C)** Using the $[water](z)$ for the new bilayer (in this case POPC) and the $\sigma_{NP}([water])$ function derived in this paper (Figure 3, Equation 4), the $\sigma_{NP}(z)$ can be determined for a given bilayer. In Panel **C**, the black line is the $\sigma_{NP}(z)$ for the DLPC bilayer used in this study, and the blue line is the $\sigma_{NP}(z)$ for a POPC bilayer. We find that the $\sigma_{NP}(z)$ for both of these bilayers are almost indistinguishable, indicating that $\sigma_{NP}(z)$ over the experimental range is not affected by changes in lipid acyl tail length or saturation.

**Figure S10. Bilayer Position Dependence of Arginine $\Delta G^{\circ}_{sc}$**

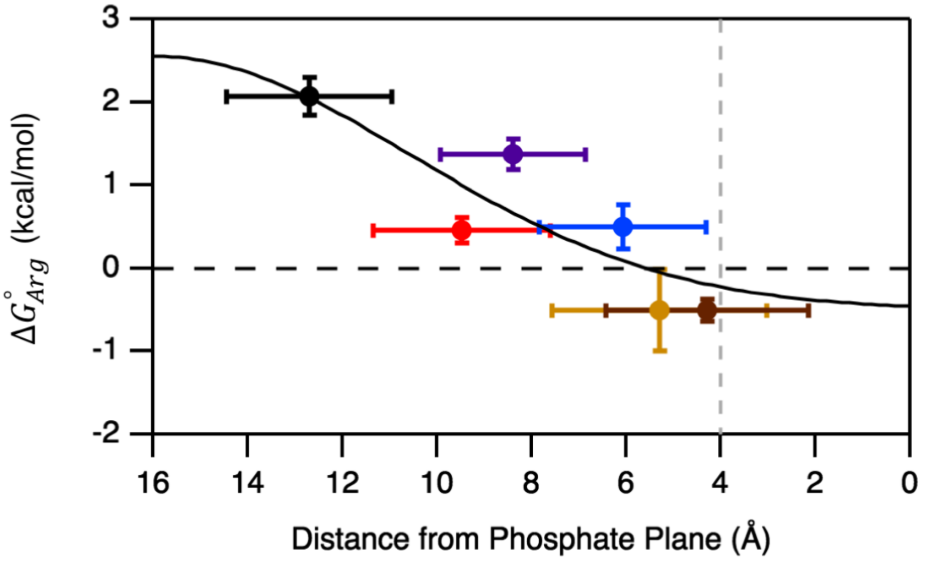

The $\Delta G^{\circ}_{sc}$ for arginine at each of the six host sites on OmpLA are plotted as a function of their position relative to the phosphate plane in the DLPC bilayer (error bars represent standard deviations for both axes). We find the energetic penalty for inserting an arginine in the dehydrated core of the bilayer is approximately 2 kcal/mol. However, it is energetically favorable to have a lipid facing arginine within 6Å of the phosphate plane. We find that $\Delta G^{\circ}_{Arg}(z)$ can be defined by described by a Gaussian distribution (black line, Table S7). The description of the position dependence of $\Delta G^{\circ}_{Arg}$ as a Gaussian function has been reported previously.[6] The vertical, gray dotted dashed line references the position of the carbonyl plane in the bilayer relative to the phosphate plane.

**Figure S11. Bilayer Position Dependence of Aromatic $\Delta G°_{sc}$**

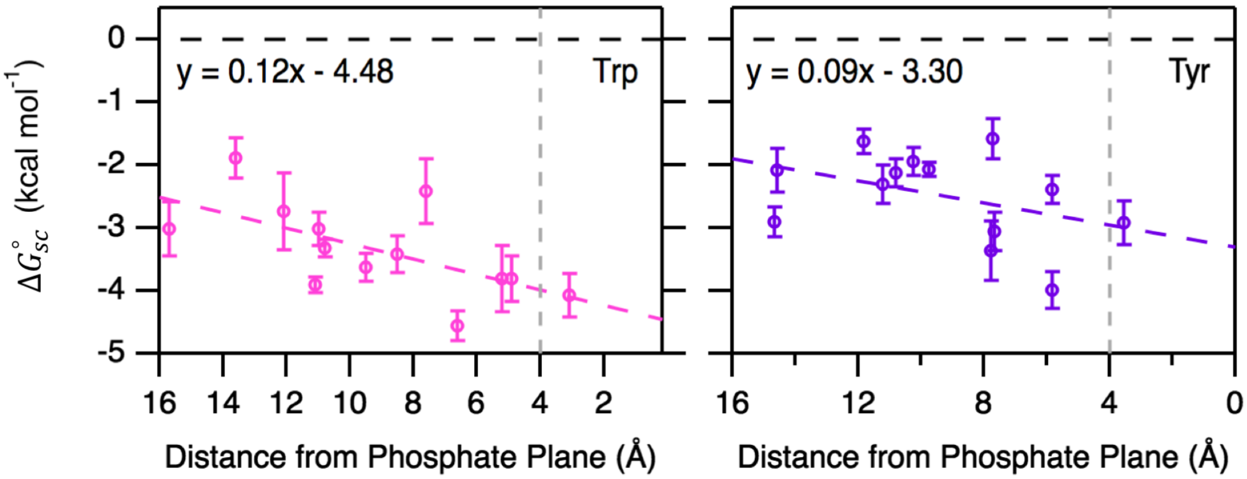

The reference-free $\Delta G^{o}_{sc}$ values for Trp and Tyr are shown as a function of bilayer position, with linear regression $\Delta G^{o}_{Trp}$ and $\Delta G^{o}_{Tyr}$ functions shown as dashed lines ($R^2$ = 0.38 and 0.20, respectively). Trp and Tyr are more favorable in the bilayer interface than the center of the bilayer, reflecting the preference of aromatic residues to be found in interfacial regions of transmembrane domains. The vertical, gray dotted dashed line references the position of the carbonyl plane in the bilayer relative to the phosphate plane.

### Supplemental Tables

#### Table S1. Average Cα positions for each OmpLA variant relative to the bilayer phosphate plane

| | Site on OmpLA | | | | | |
|---|---|---|---|---|---|---|
| | 164 | 210 | 223 | 212 | 120 | 214 |
| Ile | 6.83 ± 2.02 | 13.50 ± 1.87 | 5.98 ± 1.90 | 9.25 ± 1.72 | 11.34 ± 2.65 | 5.84 ± 2.40 |
| Leu | 7.02 ± 1.41 | 13.94 ± 1.54 | 5.93 ± 1.73 | 9.63 ± 1.86 | 12.41 ± 1.91 | 4.55 ± 1.70 |
| Met | 7.40 ± 1.71 | 12.66 ± 1.41 | 7.73 ± 1.55 | 10.3 ± 1.37 | 9.50 ± 1.80 | 5.10 ± 1.75 |
| Val | 6.76 ± 1.62 | 14.40 ± 1.34 | 6.72 ± 1.73 | 8.23 ± 1.88 | 11.92 ± 1.84 | 3.74 ± 1.91 |
| Ala | 7.43 ± 2.18 | 12.92 ± 1.82 | 5.81 ± 1.92 | 8.86 ± 1.29 | 11.24 ± 1.84 | 3.85 ± 2.93 |
| Phe | 7.68 ± 1.06 | 14.68 ± 0.86 | 8.22 ± 1.17 | 10.41 ± 1.34 | 10.13 ± 1.13 | 5.57 ± 1.25 |
| Arg | 5.30 ± 1.54 | 12.71 ± 1.54 | 6.08 ± 1.19 | 8.39 ± 1.34 | 9.47 ± 1.54 | 4.29 ± 1.69 |
| NP | 7.19 ± 1.65 | 13.68 ± 1.48 | 6.73 ± 1.60 | 9.44 ± 1.54 | 11.09 ± 1.82 | 4.78 ± 1.95 |

Individual side chain average positions and standard deviations are determined from the Gaussian fits in Figure S5. The NP row is the average position and error for all nonpolar side chains (i.e. excluding arginine), which are used as the position and error for $\sigma_{NP}$ plotted in Figure 4. All numbers are distances in angstroms from the average position of the phosphate plane in all-atom molecular dynamics simulations.

**Table S2. Stabilities and Alanine-Dependent $\Delta\Delta G°_{w,l}$ of Ile, Met, and Val OmpLA Variants**

| OmpLA Variant | $\Delta G^{o}_{w,l}$[a] (kcal mol$^{-1}$) | $\Delta\Delta G^{o}_{w,l}$[b] (kcal mol$^{-1}$) |
|---|---|---|
| 120I | -30.91 ± 0.01 | -2.72 ± 0.08 |
| 164I | -32.89 ± 0.63 | -1.27 ± 0.74 |
| 212I | -36.55 ± 0.18 | -4.12 ± 0.18 |
| 214I | -30.57 ± 0.14 | -1.61 ± 0.20 |
| 120M | -29.43 ± 0.28 | -1.24 ± 0.48 |
| 164M | -32.02 ± 0.36 | -0.40 ± 0.53 |
| 212M | -33.35 ± 0.05 | -0.92 ± 0.06 |
| 214M | -30.11 ± 0.03 | -1.15 ± 0.14 |
| 120V | -30.29 ± 0.12 | -2.10 ± 0.14 |
| 164V | -31.46 ± 0.64 | 0.16 ± 0.75 |
| 212V | -35.88 ± 0.49 | -3.45 ± 0.49 |
| 214V | -30.07 ± 0.12 | -1.11 ± 0.19 |

Errors are either standard deviations (n = 3) or standard error of the mean (n = 2)

[a]$\Delta G^{o}_{w,l}$ values are the Gibbs free energies for folding for each variant for the water ($w$)-to-lipid ($l$) transition. Chemical denaturation titrations were fit to a three state model with $m$-values held constant to previously determined values for each transition (2.03 and 7.18 kcal mol$^{-1}$ M$^{-1}$, respectively).[3] $\Delta G^{o}_{w,l}$ above are the sum of the best-fit values for the two transitions in the three-state fit.

[b]$\Delta\Delta G^{o}_{w,l}$ values were calculated by subtracting the $\Delta G^{o}_{w,l}$ for the guest variant shown above from the $\Delta G^{o}_{w,l}$ for the previously reported alanine variant at each site.[2] Error is propagated from the $\Delta G^{o}_{w,l}$ for all variants.

### Table S3. Parameters for $\sigma_{NP}$ Fits in Figure 2

| Site | Slope ($\sigma_{NP}$) | y-intercept |
|---|---|---|
| 120 | -27.3 | 0.03 |
| 164 | -18.9 | 0.20 |
| 210 | -23.8 | 0.10 |
| 212 | -24.5 | 0.03 |
| 214 | -16.1 | -0.08 |
| 223 | -22.7 | -0.12 |

The slopes and intercepts for the weighted linear regressions shown in Figure 2 are reported here.

**Table S4. Reference Free $\Delta G°_{sc}$ for Side Chains at the Six Sites on OmpLA**

| Site | Side Chain | $\Delta G^o_{sc}$ (kcal mol$^{-1}$) [a] | Site | Side Chain | $\Delta G^o_{sc}$ (kcal mol$^{-1}$)[a] |
|---|---|---|---|---|---|
| 120 | A | -1.89 ± 0.09 | 164 | A | -1.30 ± 0.41 |
| | F | -4.17 ± 0.30 | | F | -3.05 ± 0.46 |
| | I* | -3.80 ± 0.12 | | I | -2.58 ± 0.85 |
| | L | -4.05 ± 0.13 | | L | -2.51 ± 0.46 |
| | M | -3.12 ± 0.49 | | M | -1.71 ± 0.67 |
| | R | 0.46 ± 0.15 | | R | -0.50 ± 0.49 |
| | V* | -3.19 ± 0.17 | | V | -1.14 ± 0.85 |
| | W | -4.05 ± 0.13 | | W | -2.13 ± 0.51 |
| | Y | -2.46 ± 0.30 | | Y | -3.06 ± 0.47 |
| 212 | A | -1.69 ± 0.08 | 214 | A | -1.11 ± 0.13 |
| | F | -4.73 ± 0.19 | | F | -3.34 ± 0.28 |
| | I* | -3.43 ± 0.20 | | I | -2.72 ± 0.24 |
| | L | -4.31 ± 0.22 | | L | -2.32 ± 0.14 |
| | M | -2.61 ± 0.10 | | M | -2.26 ± 0.19 |
| | R | 1.37 ± 0.18 | | R | -0.50 ± 0.13 |
| | V* | -2.76 ± 0.49 | | V | -2.22 ± 0.23 |
| | W | -3.28 ± 0.14 | | W | -3.77 ± 0.36 |
| | Y | -2.05 ± 0.11 | | Y | -3.77 ± 0.29 |
| 223 | A | -1.57 ± 0.16 | | | |
| | F | -3.82 ± 0.29 | | | |
| | I | n/a | | | |
| | L | -3.40 ± 0.17 | | | |
| | M | n/a | | | |
| | R | 0.50 ± 0.26 | | | |
| | V | n/a | | | |
| | W | -4.67 ± 0.24 | | | |
| | Y | -3.04 ± 0.31 | | | |

[a]$\Delta G^o_{sc}$ are the reference-free side chain transfer free energy for a given side chain at that position on OmpLA. The $\Delta G^o_{sc}$ for alanine are calculated by multiplying the nonpolar solvation parameter determined for that site (Figure S3) by the nonpolar surface area of an alanine side chain (69.1 Å$^2$). For all other side chains, $\Delta G^o_{sc}$ is calculated by adding the $\Delta G^o_{sc}$ for alanine to the $\Delta\Delta G°_{w,l}$ measured for each side chain at that site on OmpLA (Table S1).

*$\Delta G^o_{sc}$ for beta-branched residues at sites 120 and 212 have been decreased by 0.80 and 2.38 kcal/mol respectively to reflect only the transfer free energy of these side chains and not local interactions.

**Table S5. Local water concentration and $\sigma_{NP}$ for each site on OmpLA and PagP**

| Site | $\sigma_{NP}$ (cal mol$^{-1}$ Å$^{-2}$) | Local [water] (M) |
|---|---|---|
| 210 | -23.8 | 0.15 |
| 120 | -27.3 | 0.58 |
| 111 (PagP) | -25.7 | 0.65 |
| 212 | -24.5 | 1.37 |
| 164 | -18.9 | 4.20 |
| 223 | -22.7 | 5.18 |
| 214 | -16.1 | 11.33 |

The local [water] for each site were determined using the average nonpolar Cα position listed in Table S1 and using Equation 3 (Methods section). For site 111 on PagP, analyses were carried out using previously published simulations of WT PagP[7].

**Table S6. Aromatic $\Delta G^o_{sc}$ at seven additional sites on OmpLA**

| | $\Delta G^o_{sc}$ (kcal mol$^{-1}$) [a] | | |
|---|---|---|---|
| Site | Phe | Trp | Tyr |
| 243 | -3.92 ± 0.34 | -4.07 ± 0.35 | -2.91 ± 0.34 |
| 195 | n/a | -3.62 ± 0.22 | -1.94 ± 0.22 |
| 239 | -4.74 ± 0.21 | -2.73 ± 0.61 | -2.13 ± 0.22 |
| 173 | -4.29 ± 0.20 | -3.02 ± 0.43 | -2.90 ± 0.23 |
| 162 | -3.44 ± 0.23 | -3.02 ± 0.26 | -1.63 ± 0.19 |
| 136 | -3.43 ± 0.25 | -3.41 ± 0.29 | -1.58 ± 0.32 |
| 122 | -4.27 ± 0.31 | -3.80 ± 0.52 | -2.39 ± 0.22 |

[a] $\Delta G^o_{sc}$ for these sites on OmpLA were determined using the predicted bilayer position dependent profile for $\Delta G^o_{Ala}$. The bilayer positions of each variant were previously determined using molecular dynamics simulations, and the corresponding $\Delta G^o_{Ala}$ for alanine at that position in the bilayer was used to adjust the experimentally determined $\Delta\Delta G^o_{w,l}$ at these sites by McDonald and Fleming.[2]

**Table S7. $\Delta G^{\circ}_{sc}(z)$ Equations**

| Side Chain | Function |
|---|---|
| A | $\Delta G^{\circ}_{sc}(z) = 0.069 * \left(0.87 * \left(1.0 * 10^{-6} + \left\{\frac{31.563}{1 + exp^{\frac{3.7015-z}{-1.8604}}}\right\}\right) - 25.6\right)$ |
| F | $\Delta G^{\circ}_{sc}(z) = 0.170 * \left(0.87 * \left(1.0 * 10^{-6} + \left\{\frac{31.563}{1 + exp^{\frac{3.7015-z}{-1.8604}}}\right\}\right) - 25.6\right)$ |
| I | $\Delta G^{\circ}_{sc}(z) = 0.141 * \left(0.87 * \left(1.0 * 10^{-6} + \left\{\frac{31.563}{1 + exp^{\frac{3.7015-z}{-1.8604}}}\right\}\right) - 25.6\right)$ |
| L | $\Delta G^{\circ}_{sc}(z) = 0.143 * \left(0.87 * \left(1.0 * 10^{-6} + \left\{\frac{31.563}{1 + exp^{\frac{3.7015-z}{-1.8604}}}\right\}\right) - 25.6\right)$ |
| M | $\Delta G^{\circ}_{sc}(z) = 0.118 * \left(0.87 * \left(1.0 * 10^{-6} + \left\{\frac{31.563}{1 + exp^{\frac{3.7015-z}{-1.8604}}}\right\}\right) - 25.6\right)$ |
| R | $\Delta G^{\circ}_{sc}(z) = -0.5 + 3.05exp^{\left\{-\left(\frac{z-16}{7.75}\right)^{2}\right\}}$ |
| V | $\Delta G^{\circ}_{sc}(z) = 0.116 * \left(0.87 * \left(1.0 * 10^{-6} + \left\{\frac{31.563}{1 + exp^{\frac{3.7015-z}{-1.8604}}}\right\}\right) - 25.6\right)$ |
| W | $\Delta G^{\circ}_{sc}(z) = 0.12 * z - 4.48$ |
| Y | $\Delta G^{\circ}_{sc}(z) = 0.09 * z - 3.30$ |

*z is z-position of Cα atom in the bilayer relative to the nearest phosphate plane
